# Identification of m^6^A residues at single-nucleotide resolution using eCLIP and an accessible custom analysis pipeline

**DOI:** 10.1101/2020.03.11.986174

**Authors:** Justin T. Roberts, Allison M. Porman, Aaron M. Johnson

## Abstract

Methylation at the N^6^ position of adenosine (m^6^A) is one of the most abundant RNA modifications found in eukaryotes, however accurate detection of specific m^6^A nucleotides within transcripts has been historically challenging due to m^6^A and unmodified adenosine having virtually indistinguishable chemical properties. While previous strategies such as methyl-RNA immunoprecipitation and sequencing (MeRIP-Seq) have relied on m^6^A-specific antibodies to isolate RNA fragments containing the modification, these methods do not allow for precise identification of individual m^6^A residues. More recently, modified cross-linking and immunoprecipitation (CLIP) based approaches that rely on inducing specific mutations during reverse transcription via UV crosslinking of the anti-m^6^A antibody to methylated RNA have been employed to overcome this limitation. However, the most utilized version of this approach, miCLIP, can be technically challenging to use for achieving high-complexity libraries. Here we present an improved methodology that yields high library complexity and allows for the straightforward identification of individual m^6^A residues with reliable confidence metrics. Based on enhanced CLIP (eCLIP), our m^6^A-eCLIP (meCLIP) approach couples the improvements of eCLIP with the inclusion of an input sample and an easy-to-use computational pipeline to allow for precise calling of m^6^A sites at true single nucleotide resolution. As the effort to accurately identify m^6^As in an efficient and straightforward way intensifies, this method is a valuable tool for investigators interested in unraveling the m^6^A epitranscriptome.

## Introduction

N^6^-methyladenosine (m^6^A) is a modification to RNA where a methyl group is added to the N^6^ position of adenosine. m^6^A is the most prevalent post-transcriptional modification of eukaryotic mRNA and has important roles in a variety of physiological processes including cell differentiation (Geula et al. 2015) and development (Y. Wang et al. 2014), alternative splicing (Xiao et al. 2016), and regulation of mRNA stability (X. Wang et al. 2014). m^6^A residues are typically deposited co-transcriptionally (Ke et al. 2017) onto nascent pre-mRNA molecules in the nucleus via a ‘writer’ consisting of a stable heterodimer enzyme complex composed of methyltransferase proteins METTL3/14 in association with the pre-mRNA regulator WTAP and additional accessory components such as KIAA1429 and RBM15 (Ke et al. 2017; Wu et al. 2017). This writer complex targets RNAs containing a ‘DRACH’ consensus sequence (where ‘D’ is any nucleotide but cytosine, ‘R’ is any purine, and ‘H’ is any nucleotide but guanine), with the cytosine downstream of the substrate adenine being essential for methylation (Liu et al. 2014). The human methyltransferase METTL16 can also generate m^6^A modifications, though these residues do not occur within the same ‘DRACH’ consensus motif and only a very few substrates are known (Ruszkowska et al. 2018; Doxtader et al. 2018). While m^6^A modifications have been identified throughout the transcriptome, they are most-often enriched around 3’ UTRs and stop codons (Meyer et al. 2012; Dominissini et al. 2012; Ke et al. 2015). In contrast, a similar RNA modification, N^6^, 2′-O-dimethyladenosine (m^6^Am), is located on the 5’ ends of mRNAs and is catalyzed by the methyltransferase PCIF1 (Boulias et al. 2019; Sendinc et al. 2019). Following methylation, m^6^A containing-transcripts are specifically recognized by ‘reader’ proteins, the most well characterized being members of the YTH domain family (Wu et al. 2017). Depending on the subcellular localization of these readers, recognition of m^6^A can mediate critical cellular functions. For example, YTHDC1, the main nuclear YTH protein, regulates pre-mRNA splicing (Xiao et al. 2016), nuclear export (Roundtree et al. 2017), and transcriptional repression (Patil et al. 2016), whereas binding of m^6^A via the cytoplasmic reader YTHDF2 leads to transcript decay (X. Wang et al. 2014). m^6^A residues are also dynamically reversible via ‘erasure’ by the demethylases ALKBH5 and FTO (Wu et al. 2017).

Current strategies to identify m^6^A residues are largely based on using m^6^A-specific antibodies to isolate transcripts containing methylated adenosine. The initial approaches, methyl-RNA immunoprecipitation and sequencing (MeRIP-Seq) (Meyer et al. 2012) and m^6^A-seq (Dominissini et al. 2012), involved the immunoprecipitation of ~100nt long RNA fragments bound to anti-m^6^A antibodies whereby successive sequencing and mapping of the reads results in the selective enrichment for sequences that contain m^6^A. However, as the m^6^A residue could be anywhere within the precipitated fragment, single-nucleotide resolution could only be approximated using the ‘DRACH’ motif as a guide to predict the specific m^6^A site. In contrast, more recently several techniques have demonstrated the ability to overcome this limitation by employing crosslinking and immunoprecipitation (CLIP) (Licatalosi et al. 2008) based strategies where UV light is used to crosslink the m^6^A antibody to methylated transcripts. Two of these methods, m^6^A-CLIP (Ke et al. 2015) and ‘m^6^A individual-nucleotide-resolution CLIP’ (miCLIP) (Linder et al. 2015), demonstrate that following crosslinking and IP of the antibody:RNA complex, removal of the antibody leaves a cross-linked amino acid “scar” near the m^6^A site, and reverse transcription over this scar leads to distinct mutations that arise from the increased frequency of reverse transcriptase errors at the exact nucleotide where amino acids crosslink to RNA. Specifically, the miCLIP technique showed that there is a markedly high frequency for C-to-T transitions at the obligate C that occurs one nucleotide downstream of putative m^6^A sites within the resulting cDNA (an A is incorporated into the cDNA instead of a G). These mutations can then be used to identify individual m^6^A residues via computational screening of sequencing reads (a similar method, ‘photo-crosslinking-assisted m^6^A-sequencing (PA-m^6^A-seq) (Chen et al. 2015), uses PAR-CLIP (Hafner et al. 2010) to identify m^6^A residues based on the introduction of 4-thiouridine induced T-to-C transitions near the methylated adenosine). While these strategies do resolve the lack of single-nucleotide resolution inherent in the previous non-CLIP based methods, there are several limitations that remain to be addressed. Specifically, the miCLIP protocol itself employs several steps such as radiolabeling and circularization of the cDNA library that make it technically challenging and frequently results in lower-complexity libraries. Further, while all the methods outline a strategy to identify m^6^A sites from the resulting sequencing reads, the series of steps involved requires a considerable amount of bioinformatic expertise in order to obtain an actual set of m^6^A positions.

We have developed an updated antibody-based approach to accurately call m^6^A residues at single-nucleotide resolution. m^6^A-eCLIP (meCLIP) is a modification of the existing eCLIP protocol (Van Nostrand et al. 2016) with changes specifically designed to identify m^6^As. Compared to existing strategies, the protocol is technically simplified and includes a comprehensive computational pipeline that runs to completion once it is executed. We have successfully validated this strategy on several cell lines and confirmed its ability to accurately call individual m^6^A residues throughout the transcriptome in a high-throughput and more straightforward manner compared to previous approaches.

## Results

### Overview of eCLIP Library Preparation

Our meCLIP approach utilizes UV crosslinking to covalently link an anti-m^6^A antibody to fragmented polyA-selected transcripts containing m^6^A and then immunoprecipitates the antibody-bound RNA. This antibody:RNA complex is then ran on an SDS-PAGE gel and transferred to a nitrocellulose membrane to remove any non-crosslinked RNA. Following treatment with Proteinase K to remove nearly all of the antibody except the crosslinked amino acid, the RNA is isolated, one adapter is ligated, and the RNA is converted into cDNA. All first-strand cDNA products receive a second adapter required for sequencing, ensuring that efficiency of the reverse transcriptase crossing the amino acid “scar” does not impede the library preparation. Reverse transcription over the anti-m^6^A crosslink site results in C-to-T mutations in the template strand read from the resulting sequencing, and a custom algorithm then identifies sites of elevated C-to-T conversion frequency that occur within the m^6^A consensus motif. An overview of the library preparation can be seen in Figure 1A.

**Figure 1.**
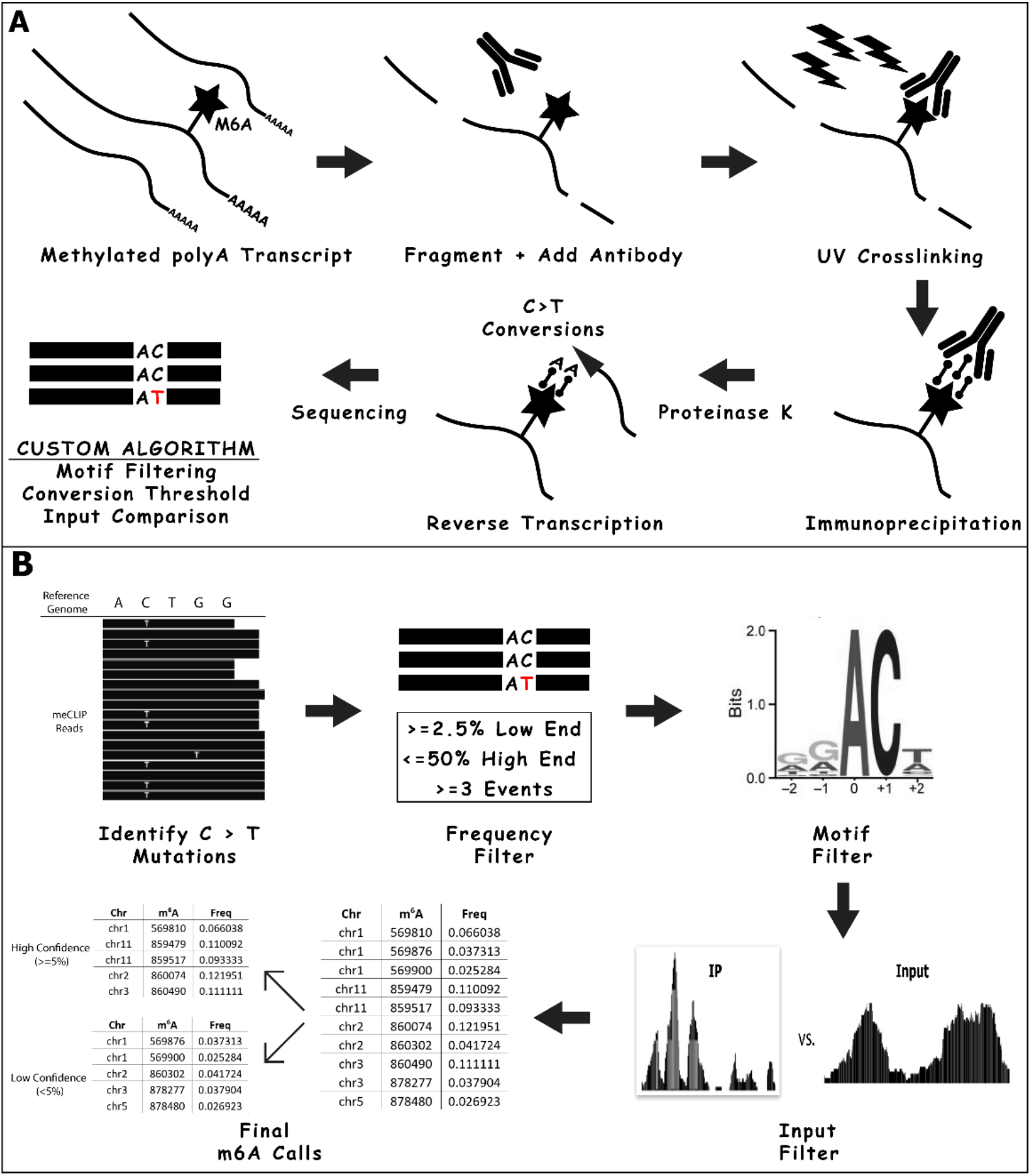
Overview of the meCLIP strategy, including summary of library preparation and the subsequent algorithm to identify m^6^A residues from the sequencing reads. A) Following isolation of mRNA from total RNA samples, the transcripts are fragmented and UV crosslinked to anti-m^6^A antibody (top). Following immunoprecipitation (bottom right), the antibody is removed and the RNA is reverse transcribed. Residual amino acid adducts resulting from the RNA:antibody crosslinking cause C-to-T mutations that are detectable in the resulting sequencing reads (bottom middle). These mutations are used as input for a custom algorithm that identifies sites of elevated C-to-T conversion frequency that occur within the m^6^A consensus motif (bottom left). B) Following sequencing, the resulting reads are used for a custom algorithm that uses the ‘mpileup’ command of SAMtools (Li et al. 2009) to identify sites of elevated C-to-T mutations. These positions are then filtered based on the frequency of the conversion (>=2.5% and <=50% with a minimum of 3 events) and their occurrence within the m^6^A consensus motif (‘RAC’, where ‘R’ is any purine). Finally, the filtered positions are compared to the similarly analyzed input sample and any overlapping positions are removed. The resulting m^6^A calls are categorized into low and high confidence sets based on the mutational frequency (<5% for low, >=5% for high).

### Optimization of RNA Fragmentation Improves Successful Library Generation

We have found that initial optimization of the cation-based RNA fragmentation step can increase the quality of the final library. While too little fragmentation results in amplicons that are outside the recommendations for Illumina sequencing (200-500bp for NovaSEQ 6000), over-fragmenting can result in a severe reduction in yield of appropriately sized immunoprecipitated RNA for input into the library preparation steps. While the actual conditions for fragmentation depend on the individual RNA sample, important factors to consider include: 1) the concentration of input RNA; 2) the duration of time and temperature that the RNA is fragmented; and 3) the length of reads used in sequencing. Based on these points, we recommend that a trial with total RNA be fragmented at 70°C with various durations ranging from 3 to 15 minutes and then analyzed on a TapeStation (Agilent) using High Sensitivity RNA ScreenTape (if unavailable, visualization by agarose gel electrophoresis can be performed instead). Further adjustments can then be made for the actual polyA RNA sample taking into consideration that polyA selected RNA tends to fragment slightly faster. We recommend an optimal fragment size of 100 to 200nt (for sequencing on NovaSeq with 2×150 run). For reference, we have included sample TapeStation results from appropriate and undesirable fragmentations (Supplemental Figure 1). The TapeStation coupled with High Sensitivity RNA ScreenTape requires minimal RNA material for analysis allowing for the same sample to be analyzed and then subsequently used in the experiment.

### Improved Adapter Ligation to Increase Efficiency

Instead of addition of both sequencing adapters to the original RNA fragments, as in the original HITS-CLIP protocol (Licatalosi et al. 2008), or using circular ligation as implemented in iCLIP, which is challenging to perform efficiently, the eCLIP protocol (Van Nostrand et al. 2016) adds adapters for sequencing in two separate steps. The first step uses an indexed 3’ RNA adapter that is ligated to crosslinked RNA fragment while still on the immunoprecipitation beads, and the second is a 3’ ssDNA adapter that is ligated to cDNA following reverse transcription. The first 3’ RNA adapter is ‘in-line-barcoded’ and may consist of a number of matched combinations (A01+B06, X1A+X1B, etc.) that are detailed in the original eCLIP protocol and included here as well. The second ssDNA adapter (rand3Tr3) contains a unique molecular identifier (UMI) that allows for determination of whether two identical sequenced reads indicate two unique RNA fragments or PCR duplicates of the same RNA fragment. Therefore, the resulting reads generally have the following structure:

Read 1 – NNNNNGCTATT [Sequenced Fragment (RC)] NNNNNNNNNNAGATCGGAAGAGCAC
Read 2 – NNNNNNNNNN [Sequenced Fragment] AATAGCANNNNN

where read 1 begins with the barcoded RNA adapter (X1A displayed here) and read 2 (corresponding to the sense strand) begins with the UMI (either N5 or N10) followed by a sequence corresponding to the 5′ end of the original RNA fragment or reverse transcription termination site. To account for errors in the UMI itself (which would impact the accuracy of quantifying unique molecules at a given genomic locus), our downstream analysis pipeline uses the software package UMI-tools (Smith, Heger, and Sudbery 2017) which employs network-based algorithms to correctly identify true PCR duplicates.

### Standardized Strategy to Reduce PCR Duplication

In addition to not requiring any radiolabeling, compared to iCLIP, the eCLIP protocol also decreases the required amplification by up to ~1,000-fold resulting in far less reads being removed due to PCR duplication of the same molecule. The original protocol recommends performing qPCR on a diluted sample of cDNA and adjusting the final library amplification based on the dilution (i.e. 3 less cycles than the 1:10 dilution). Our approach takes this recommendation a step further by performing amplification on diluted cDNA across a range of cycles and running them on a polyacrylamide gel to allow for visualization of the optimal amount of amplification (Supplemental Figure 2). This extra step ensures that the final library will not be saturated with PCR duplicates while still allowing for adequate sequencing depth to identify m^6^As.

### Straightforward Analysis Pipeline Using Snakemake

The current most widely used m^6^A site identification strategy (miCLIP) relies on ‘crosslinking-induced mutation site analysis, CIMS (now a part of the CLIP Tool Kit (CTK) software package (Shah et al. 2017)) which was originally designed to identify sites of RNA:protein crosslinking from CLIP data. While this method does ultimately provide a set of m^6^A sites deduced from the C-to-T mutations in the sequencing reads, the implementation requires considerable bioinformatic expertise in the form of manually installing prerequisite software and executing the individual tools from the command line step-by-step. In comparison, our approach implements the workflow management system Snakemake (Köster and Rahmann 2012) to streamline the process, requiring only a single configuration file and no manual installation of software libraries. Specifically, as opposed to running multiple scripts one at a time, Snakemake workflows combine the execution of all the component commands into a human readable file that is easily modifiable.

Once the workflow is executed, the reads are first assessed for appropriate quality and presence of adapters which are removed accordingly. The reads are then mapped to RepBase (Bao, Kojima, and Kohany 2015) to remove repetitive elements and ribosomal RNA and then to the actual reference genome itself. Following mapping, any PCR duplicates are collapsed within the alignment file which is then sorted and indexed in preparation for calling mutations. The output files of each of these steps is automatically compiled and presented as a summary file to the user for reference. Finally, the custom algorithm (outlined in Figure 1B) identifies variations from the reference genome at each position and then specifically identifies C-to-T conversions occurring between a frequency threshold of greater than or equal to 2.5% and less than or equal to 50%. Those positions meeting these thresholds are then analyzed for the presence of the m^6^A consensus motif ‘RAC’ (where R is any purine). After comparing to the corresponding input sample (described below), a list of m^6^A sites are reported in the form of their individual coordinates within the genome along with gene and transcript annotations and supporting C-to-T mutation frequency. A metagene profile of the identified residues summarizing how they localize within transcripts is also automatically generated (Figure 2). All these steps are executed automatically with minimal input required from the user, taking approximately 6-8 hours depending on the size of the sequencing library and available computational resources. The pipeline itself can be scaled seamlessly to clusters and cloud servers depending on the available user environment without the need to modify the workflow.

**Figure 2.**
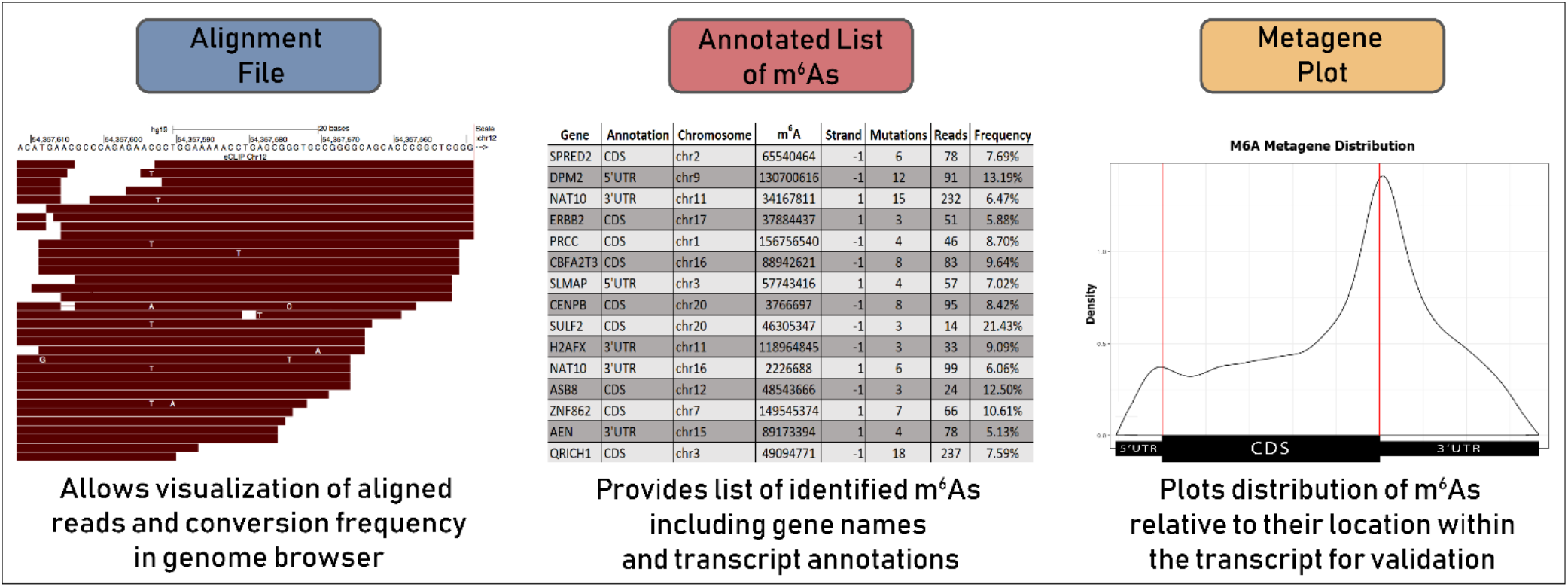
Overview of output files provided by the workflow. In addition to a summary file consisting of the relevant outputs and logs from the prerequisite software used in the workflow (not shown), an alignment file (BAM) consisting of reads that were successfully mapped to the genome is supplied for visualization of overall library quality and C-to-T conversion frequencies relative to identified m^6^A sites. A tab-delimited list of called m^6^As (sorted by confidence) with their genomic location, gene and transcript annotation, number of supporting C-to-T mutations and relative conversion frequency are also provided (two separate BED files containing just the genomic coordinates of m^6^A residues in each confidence category are also generated for use in downstream analysis / visualization). A metagene profile summarizing where in the transcript the identified m^6^A residues occur is also automatically generated using MetaPlotR (Olarerin-George and Jaffrey 2017).

### Use of Input Sample to Control for Conversion Calls Not Arising From m^6^A Antibody

Our method includes a corresponding input sample that is used to identify C-to-T mutations that occur in the absence of the anti-m^6^A antibody. A small aliquot of fragmented RNA is taken prior to introduction of the antibody and subsequent crosslinking (representing ~5% of the total sample) and prepared concurrently with the immunoprecipitated sample after the antibody is removed. Following sequencing, the same m^6^A identification workflow is performed on the input reads which identifies C-to-T mutations that are not induced from anti-m^6^A antibody. The computational pipeline automatically takes these ‘false m^6^As’ and compares them to the list of m^6^A sites obtained from the true m^6^A immunoprecipitation. By removing any positions that occur in both sets, our strategy specifically identifies adenosine residues that are recognized by the anti-m^6^A antibody. Our experiments to date have identified that such contaminating mutations account for 5-10% of the initial residues called from the immunoprecipitation. As these m^6^A residues would otherwise likely remain included in other strategies, we consider this step to be a significant improvement and ultimately required for confidence in accurate identification of m^6^As. Further, as the input essentially represents an RNA-seq experiment of the same sample, it can be used to gauge overall gene expression within the sample.

### Successful Identification of m^6^A Sites Categorized by Confidence

Using the meCLIP strategy outlined above, we have successfully identified over 50,000 m^6^A residues in MCF-7 and MDA-MB-231 breast cancer cell lines. We find a considerable difference in the number of m^6^A sites called between experimental replicates of the same cell line (Table 1). Whether this is a result of variability in crosslinking efficiency, variable frequency of reverse transcription errors, or evidence of the dynamic nature of m^6^A deposition itself, these results indicate that calling m^6^As on replicates of the same sample multiple times and taking the consensus is often the best strategy to be confident in specific modifications. We also noted that the consensus among replicates increases when the C-to-T mutation frequency at a given residue is above 5%, and that the majority of conversions are not located within the m^6^A ‘RAC’ consensus motif when the mutation frequency is less than 5% (Supplemental Figure 3). Binning all sites between 1-50% mutation frequency into bins of 2.5%, we observed that the 2.5-4.9% bin has an ‘RAC’ motif occurrence well below the ~50% which occurs in the next higher bin. A higher mutation rate is observed across the 5-50% range. When the mutation frequencies are 50% and above, the motif occurrence trends close to 12.5% frequency of random chance of having ‘RA’ upstream of the C. Based on these observations, we therefore have chosen to automatically categorize our m^6^A site calls into low and high confidence based on the frequency of the C-to-T mutation at that residue where low confidence is 2.5-5% and high confidence is 5-50%).

**Table 1.** Summary of m^6^A residues identified in two breast cancer cell lines using meCLIP. The overlap in site calls between replicates was >50% of Replicate #2 for both cell lines tested. The total consensus is higher than within confidence categories, indicating that many called m^6^A sites differ between the replicates in which confidence category they reside in. The consensus between replicates was significantly lower in the ‘Low Confidence’ category compared to ‘High Confidence’.

**MCF-7**
| **Experiment** | **Total m<sup>6</sup>A** | **Low Confidence (<5%)** | **High Confidence (>=5%)** |
| --- | --- | --- | --- |
| *Replicate #1* | 22,139 | 6,908 | 15,231 |
| *Replicate #2* | 12,827 | 6,788 | 6,039 |
| **Consensus** | **6,679** | **1,077** | **2,720** |

**MDA-MB-231**
| **Experiment** | **Total m<sup>6</sup>A** | **Low Confidence (<5%)** | **High Confidence (>=5%)** |
| --- | --- | --- | --- |
| *Replicate #1* | 31,198 | 12,205 | 18,993 |
| *Replicate #2* | 15,477 | 5,255 | 10,222 |
| **Consensus** | **9,861** | **1,569** | **3,950** |

### Comparison to Sites Called Using Previous Strategies

We performed meCLIP on HEK-293 cells and compared the identified m^6^As with those reported in the original miCLIP paper (Linder et al. 2015). We found 4,172 m^6^A sites, of which 1,307 (31.3%) were also called using miCLIP. We observed that the miCLIP approach identified significantly more residues compared to meCLIP (Supplemental Figure 4). To further investigate this difference, we analyzed our breast cancer eCLIP reads (Table 1) using the miCLIP analysis pipeline. miCLIP typically employs two different methods, crosslinking-induced truncations and mutations, i.e. CITS and CIMS, for calling m^6^A sites from CLIP reads, however given that our library preparation does not often result in truncations we only compared the number of residues reported from CIMS. While initial analyses found that miCLIP identified roughly twice as many residues compared to our strategy, we noted that there are two main differences between the m^6^A calling methods. First, miCLIP only requires a C-to-T conversion frequency of 1% with a minimum of 2 reads supporting the mutation to call the m^6^A site (for example, only 2 conversions within 200 reads). Second, unlike our strategy, the miCLIP method does not require the identified residue to occur within the m^6^A consensus motif. We found that when we compared the two analytical approaches using our thresholds the total number of m^6^A sites called between them was roughly equivalent (Figure 3), with the remaining variability in identified residues likely due to differences in the library analysis (our strategy uses the splicing-aware RNA aligner STAR (Dobin et al. 2013) to map reads whereas miCLIP uses NovoAlign/bwa (Li and Durbin 2009)). Overall, the increased thresholding in our meCLIP approach produces a set of m^6^A site calls with confidence metrics that are useful for subsequent follow-up of specific modification events.

**Figure 3.**
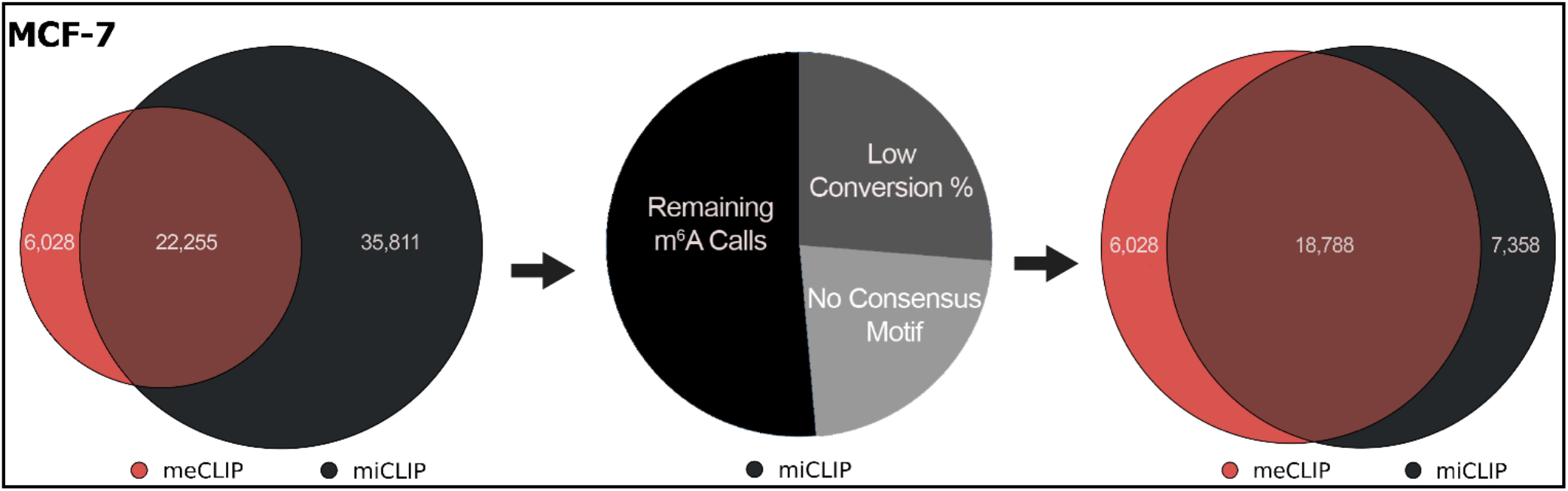
Comparison of m^6^A sites identified using the meCLIP analysis pipeline versus those identified from the miCLIP pipeline. The leftmost Venn diagram depicts the original raw numbers of m^6^As identified from both strategies from the same raw data. The pie chart in the middle shows the breakdown of the 58,066 m^6^As reported from miCLIP based on whether the residue is supported by a low conversion frequency (<2.5%) and if it occurs within the m^6^A ‘RAC’ consensus motif. The Venn diagram on the right illustrates how the m^6^As correlate between the two strategies after miCLIP residues that did not meet the threshold of our identification strategy (3 mutations, >=2.5% conversion frequency, and occurrence within the consensus motif) were removed.

### Experimental Validation of m^6^A Sites via RNA Immunoprecipitation

To experimentally confirm that the identified m^6^As are present within transcripts, we performed RNA immunoprecipitation followed by RT-qPCR on a select number of residues called in MCF-7 cells. These individual m^6^A sites were chosen based on where in the gene they occurred (i.e. 5’ UTR, 3’UTR, etc.) in order to validate the ability of our method to identify m^6^A residues across a wide spectrum of locations. For all the profiled m^6^As, we saw a significant increase in the amount of RNA recovered using anti-m^6^A antibody compared to IgG controls (Figure 4).

**Figure 4.**
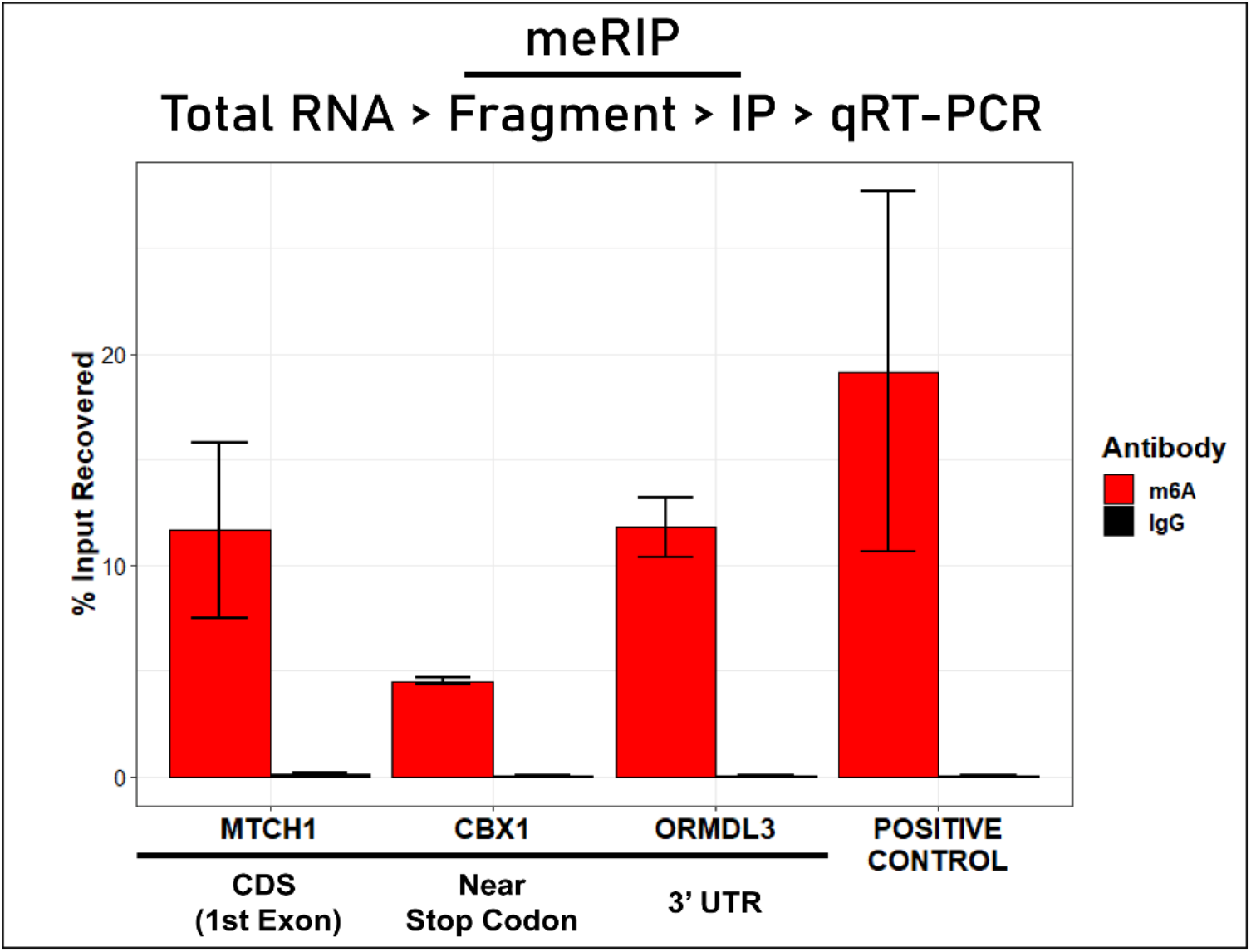
Experimental validation of select m^6^A residues. RNA immunoprecipitation using anti-m^6^A antibody (meRIP) followed by RT-qPCR was used (protocol outlined in top flowchart) to confirm the presence of m^6^A within the transcripts. Residues were chosen based on their location within the gene in order to gauge the ability of our method to identify m^6^A sites over a diverse profile of positions. Enrichment is measured as the percent of input recovered from the immunoprecipitation with anti-m^6^A compared to amount of input recovered using anti-IgG control. ‘Positive Control’ is a known m^6^A site within the EEF1A1 gene.

## Discussion

Research into RNA modifications over the past decade has shown that m^6^A is involved in most aspects of RNA biology (Zhao, Roundtree, and He 2016; Gilbert, Bell, and Schaening 2016). This ubiquitous regulation of cellular processes underscores the need to accurately and reliably identify m^6^A residues in a high-throughput transcriptome-wide manner so that context-specific m^6^A function can be better understood. While the use of antibody-based methods to identify specific m^6^A residues has become commonplace, these strategies are still challenging. We have outlined several improvement steps in our protocol which help generate m^6^A site lists that can guide subsequent research into specifically modified RNAs. These include optimization of the RNA fragmentation step, utilization of tools that account for errors in the UMI, and employment of strategies that further reduce PCR duplication. Notably, we have also streamlined the downstream analysis steps by implementing a workflow management system that automates the process of calling m^6^As.

While our meCLIP approach to identify m^6^A residues does offer clear advantages compared to previous strategies, a frequently cited challenge for using current m^6^A identification protocols is the large amount of input mRNA required for effective immunoprecipitation and sequencing. Our experience with eCLIP suggests that the input amounts we describe for our protocol could be reduced while still producing quality results, although we have not yet systematically tested the range of adequate RNA input. Consistent with recent reports (Zeng et al. 2018), however, we find that the number of unique m^6^A sites identified does increase with higher amounts of input RNA. Therefore, in addition to ensuring accurate quantification of starting material via the methods outlined in our protocol, we also highly encourage the use of multiple replicates when starting RNA material is limited to gain confidence in the identified m^6^A residues. Further, while the majority of m^6^A deposition in humans occurs via the METTL3/METTL14 writer complex, a subset is generated via another methyltransferase, METTL16 (Ruszkowska et al. 2018; Doxtader et al. 2018). In contrast to METTL3/METTL14-dependent m^6^As which are typically found within a ‘DRACH’ motif and often near stop codons, m^6^A modifications generated by METTL16 do not occur within a defined sequence motif and are more often found in introns and within noncoding RNAs (Ruszkowska et al. 2018). As our m^6^A identification protocol uses the consensus motif as a filtering mechanism and also specifically isolates mRNA via poly(A) selection, identification of METTL16-dependent m^6^As will therefore likely be limited. Similarly, although our anti-m^6^A antibody also recognizes the related RNA modification m^6^Am located on the 5’ ends of mRNAs, m^6^Am is not invariably followed by a cytidine (Boulias et al. 2019) and thus could be filtered out in our pipeline as well. However, previous reports have noted that antibody-induced A-to-T transitions at the m^6^Am site itself are also frequently observed. If specific modifications such as METTL16-dependent m^6^A and m^6^Am are of interest, the motif filtering step and conversion event of interest can be easily modified within the Snakemake workflow. The ease of making such *ad hoc* changes to the pipeline coupled with the automatic generation of datasets makes such focused identification approaches feasible and illustrates yet another benefit of our method.

In a further effort to overcome some of the limitations of previous m^6^A identification methods, recently several antibody-independent strategies have been developed to identify sites of m^6^A modifications. DART-seq (Meyer 2019) (deamination adjacent to RNA modification targets) uses a chimeric protein consisting of the YTH ‘m^6^A-reader’ domain fused to the cytidine deaminase APOBEC1 in cells to induce C-to-U deamination events at sites adjacent to m^6^A residues and then detects the mutations using standard RNA-seq. Notably, the DART-seq method only calls for as low as 10 ng of total RNA. Another pair of methods, MAZTER-seq (Garcia-Campos et al. 2019) and m^6^A-REF-seq (Kong et al. 2019), utilize the ability of the RNA endonuclease MazF to cleave single-stranded RNA immediately upstream of unmethylated sites occurring in ‘ACA’ motifs, but not within their methylated ‘m^6^A-CA’ counterparts (Imanishi et al. 2017). Finally, m^6^A identification approaches utilizing Oxford Nanopore’s direct RNA sequencing technology, including the MINES method (Lorenz et al. 2019; Price et al. 2019), have been developed to further facilitate accurate m^6^A detection. While all these methods offer unique advantages, they are not without limitations. For instance, the RNase MazF-based methods only allow for identification of the subset of m^6^As occurring within the defined motif ‘ACA’ (estimated at ~16-25% of methylation sites (Garcia-Campos et al. 2019)), making it more of a complementary strategy to quantify and validate select m^6^A residues rather than a standalone identification approach. The most significant barrier to such methods’ widespread utility however is the lack of a dedicated and straightforward analysis pipeline. We feel that coupling our m^6^A identification protocol with a package manager to easily install software dependencies and a workflow engine that automates the execution of each script is extremely valuable to those researchers with limited bioinformatic expertise and consider such an inclusion one of the most notable advantages to our calling method.

In summary, we have significantly improved the m^6^A CLIP library preparation to increase library complexity and introduced confidence metrics in identified m^6^A residues. We have also incorporated an easy-to-use analysis pipeline to facilitate the straightforward generation of lists and relevant figures detailing m^6^A deposition. Taken together, we believe our meCLIP approach to identify m^6^A modifications offers powerful benefits to investigators interested in deciphering the intricacies of m^6^A biology.

## Methods

### RNA Isolation and Fragmentation Assay

Unless indicated otherwise, the HEK-293 and MDA-MB-231 cells that were used for the described experiments contain a transgene that overexpresses the lncRNA HOTAIR. Cells were cultured in appropriate media and total RNA was isolated using the Trizol (15596018, Invitrogen) method until ~1mg of total RNA was obtained (for MCF-7 cells this was eight to ten 15cm cell culture plates at ~80-100% confluency). The total RNA samples were combined and diluted to make a 1 μg/μL stock solution (saving 1uL to assess quality of RNA on TapeStation). To determine the optimal duration of fragmentation for the desired size (100-200 nucleotides in length), a small amount (i.e. 2μg) of total RNA was fragmented using RNA Fragmentation Reagents (AM8740, Ambion) for times ranging from 3 to 15 minutes. Following the manufacturer’s protocol, combining the appropriate amount of RNA (2μL) with nuclease free water to the recommended reaction volume of 9μL, adding 1μL 10x Fragmentation Reagent, incubating at 70°C for designated time, and then immediately quenching with 1μL Stop Reagent and placing on ice. The fragment size produced from each time point is then visualized by running a heavily diluted sample (~3 ng/μL) on an Agilent TapeStation 4200 with High Sensitivity RNA Screen Tape, and the appropriate fragmentation time for the actual poly(A) sample can be approximated based on these results (see Supplemental Figure 1).

### Poly(A) Selection and Fragmentation

Poly(A) selection was performed using the Magnosphere^®^ Ultrapure mRNA Purification Kit (9186, Takara) where 100μL of the magnetic beads were combined with 250μL of total RNA (1 μg/μL) and the beads were reused four times following the manufacturer’s protocol (the volume of binding buffer that the beads are resuspended in was scaled up to match the volume of RNA, i.e. 250μL). The isolated poly(A) RNA was collected in four 50uL aliquots, combined, and 1μL was used to assess the quality of mRNA selection and determine percent recovery / concentration via TapeStation (the recommended amount of input mRNA is 5 to 20μg). The RNA sample was then ethanol precipitated overnight at −20C using standard methods with GlycoBlue Coprecipitant (AM9515, Invitrogen) added to the solution for easier recovery. The precipitated poly(A) RNA was resuspended in 15μL of nuclease free water and fragmented at 70°C for the amount of time determined previously (to further optimize the duration, initially 2μg of the polyA sample was fragmented and visualized on the TapeStation as described above; adjustments were then be made based on the observed size, or just repeated with 2μg aliquots if the size is appropriate). A small amount (~500ng) of fragmented RNA was saved for use as the input sample (see ‘Input Sample Preparation’).

### Crosslinking and Immunoprecipitation of m^6^A Containing Transcripts

The remaining fragmented RNA was resuspended in 500μL Binding/Low Salt Buffer (50mM Tris-HCl pH 7.4, 150mM Sodium chloride, 0.5% NP-40), then 2μL RNase Inhibitor (M0314, NEB) and 10μL (1 mg/ml) m^6^A antibody (Abcam Ab151230) were added and the sample was incubated on a rotator for 2 hours at 4°C. The RNA:antibody sample was transferred to one well of a pre-chilled 12-well plate and crosslinked twice at 150 mJ/cm^2^ (254nm wavelength) using a Stratalinker UV Crosslinker. 50μL of Protein A/G Magnetic Beads (88803, Pierce) were aliquoted into a fresh tube and washed twice with 500μL Binding/Low Salt Buffer. The beads were resuspended in 100μL Binding/Low Salt Buffer, added to the crosslinked RNA:antibody sample and the bead mixture was incubated at 4°C overnight with rotation. The next day, the beads were washed twice with 900μL High Salt Wash Buffer (50mM Tris-HCl pH 7.4, 1M Sodium chloride, 1mM EDTA, 1% NP-40, 0.5% Sodium deoxycholate, 0.1% Sodium Dodecyl Sulfate) and once with 500μL Wash Buffer (20 mM Tris-HCl pH 7.4, 10 mM Magnesium chloride, 0.2% Tween-20). The beads were resuspended in 500μL Wash Buffer.

### eCLIP-seq Library Preparation

#### FastAP Treatment

The beads were magnetically separated, removed from the magnet, and the supernatant was combined with 500μL 1x Fast AP Buffer (10mM Tris pH 7.5, 5mM Magnesium Chloride, 100mM Potassium Chloride, 0.02% Triton X-100). The beads were then placed on the magnet for 1 minute and the combined supernatant was removed. The beads were washed once with 500μL 1x Fast AP Buffer (following this wash, the input sample can be prepared concurrently following the ‘Input Sample Preparation’ instructions below). The beads were resuspended in Fast AP Master Mix (79μL nuclease free water, 10μL 10x Fast AP Buffer, 2μL RNase Inhibitor, 1μL TURBO DNase (AM2238, Invitrogen), 8μL Fast AP Enzyme (EF0654, Thermo Scientific)) was added and the sample was incubated at 37°C for 15 minutes with shaking at 1200 rpm.

#### PNK Treatment

PNK Master Mix (224μL nuclease free water, 60μL 5x PNK Buffer (350mM Tris-HCl pH 6.5, 50mM Magnesium Chloride), 7μL T4 PNK (EK0031, Thermo Scientific), 5μL RNase Inhibitor, 3μL Dithiothreitol (0.1M), 1μL TURBO DNase) was added to the sample and then incubated at 37°C for another 20 minutes with shaking. The beads were washed once with cold 500μL Wash Buffer, once with cold 500μL Wash Buffer and then cold 500μL High Salt Wash Buffer combined in equal volumes, once with cold 500μl High Salt Wash Buffer and then cold 500μL Wash Buffer combined in equal volumes, and once again with cold 500μL Wash Buffer. The beads were resuspended in 500μL Wash Buffer.

#### 3’ RNA Adapter Ligation

The beads were magnetically separated, removed from the magnet and then 300μL 1x RNA Ligase Buffer (50mM Tris pH 7.5, 10mM Magnesium Chloride) was added to the supernatant. The beads were placed on a magnet for 1 minute and then the combined supernatant was removed. The beads were washed twice with 300μL 1x RNA Ligase Buffer and then resuspended in 3’ RNA Ligase Master Mix (9μL nuclease free water, 9μL 50% PEG 8000, 3μL 10x RNA Ligase Buffer (500mM Tris pH 7.5. 100mM Magnesium Chloride), 2.5μL T4 RNA Ligase I (M0437, NEB), 0.8μL 100% DMSO, 0.4μL RNase Inhibitor, 0.3μL ATP (1mM), and 2.5μL (2μM) each of two matched barcoded RNA adaptors (X1A and X1B). The sample was incubated at room temperature for 75 minutes with flicking every 10 minutes.

#### Gel Loading / Membrane Transfer

The beads were washed once with cold 500μL Wash Buffer, once with cold 500μL Wash Buffer and then cold 500μL High Salt Wash Buffer combined in equal volumes, once with cold 500μL High Salt Wash Buffer, once with equal volumes of cold 500μL High Salt Wash Buffer and then 500μL Wash Buffer, and once with cold 500μL Wash Buffer. The beads were resuspended in 20μL Wash Buffer and 7.5μL 4x NuPAGE LDS Sample Buffer (NP0007, Invitrogen) and 3μL DTT (0.1M) was added. The sample was incubated at 70°C for 10 minutes with shaking at 1200 rpm and then cooled on ice for 1 minute. The beads were magnetically separated and the supernatant was transferred to a new tube. The sample was run on ice at 150V in 1x MOPS Buffer for 75 minutes on a Novex NuPAGE 4-12% Bis-Tris Gel (NP0321, Invitrogen) with 10μL dilute protein ladder (12μL Wash Buffer, 4μL M-XStable Protein Ladder (L2011, UBPBio), 4μL NuPAGE Buffer) on each side. The sample was then transferred to a nitrocellulose membrane overnight on ice at 30V. After transfer, the area of the membrane containing the antibody:RNA sample (between 20 kDa and 175 kDa) was cut and sliced into small (~2mm) pieces and placed into an Eppendorf tube.

#### Antibody Removal / RNA Cleanup

The membrane slices were incubated with 40μL Proteinase K (3115828001, Roche) in 160μL PK Buffer (100mM Tris-HCl pH 7.4, 50mM Sodium Chloride, 10mM EDTA) at 37°C for 20 minutes with shaking at 1200 rpm. An equal volume of PK Buffer containing 7M Urea was added to samples and incubated at 37°C for 20 minutes with shaking at 1200rpm, then 540μL Phenol:Chloroform:Isoamyl Alcohol (25:24:1) was added and incubated at 37°C for another 5 minutes with shaking at 1200rpm. The sample was centrifuged for 3 minutes at max speed and the aqueous layer was transferred to a 15mL conical. The RNA was isolated using the RNA Clean & Concentrator-5 Kit (R1013, Zymo) according to manufacturer’s instructions.

#### Input Sample Preparation

The input sample (~1μL) was combined with FastAP Master Mix (19μL nuclease free water, 2.5μL 10x Fast AP Buffer, 2.5μL Fast AP Enzyme, 0.5μL RNase Inhibitor) and incubated at 37°C for 15 minutes with shaking at 1200 rpm. PNK Master Mix (45μL nuclease free water, 20μL 5x PNK Buffer, 7μL T4 PNK, 1μL RNase Inhibitor, 1μL Dithiothreitol (0.1M), 1μL TURBO DNase) was added to the sample and then incubated at 37°C for another 20 minutes with shaking. The RNA sample was isolated using Dynabeads MyONE Silane (37002D, ThermoFisher Scientific). Briefly, 20μL of beads were magnetically separated and washed once with 900μL RLT Buffer (79216, Qiagen), resuspended in 300μL RLT Buffer, and added to the sample. The bead mixture was combined with 615μL 100% ethanol and 10μL sodium chloride (5M), pipette mixed, and incubated at room temperature for 15 minutes on a rotor. The sample was placed on a magnet, the supernatant was removed, and then resuspended in 1mL 75% ethanol and transferred to a new tube. After 30 seconds the bead mixture was placed on a magnet, the supernatant was removed, and the sample was washed twice with 1mL 75% ethanol, waiting 30 seconds between each magnetic separation. After the final wash, the beads were air dried for 5 minutes, resuspended in 10μL nuclease free water and incubated at room temperature for 5 minutes. The sample was magnetically separated and the elution was transferred to a new tube (an aliquot of this elution can be taken and stored at −80°C for backup if desired). The remaining eluted sample was combined with 1.5μL 100% DMSO, 0.5μL RiL19 RNA adapter, incubated at 65°C for 2 minutes, and placed on ice for 1 minute. The sample was then combined with 3’ RNA Ligase Master Mix (8μL 50% PEG 8000, 2μL 10x T4 RNA Ligase Buffer (B0216L, NEB), 1.5μL nuclease free water, 1.3μL T4 RNA Ligase I (M0437, NEB), 0.3μL 100% DMSO, 0.2μL RNase Inhibitor, 0.2μL ATP (1mM)) and incubated at room temperature for 75 minutes with mixing by flicking the tube every ~15 minutes. The sample was then re-isolated using Dynabeads MyONE Silane following the same procedure described above (after the beads were initially washed with RLT Buffer, the sample was resuspended in 61.6μL RLT Buffer instead of 300μL and an equal volume of 100% ethanol was added). The eluted input sample was then prepared simultaneously with the immunoprecipitated (IP) sample following the same instructions.

#### Reverse Transcription / cDNA Clean Up

The samples (IP and input) were reverse transcribed using the oligonucleotide AR17 and SuperScript IV Reverse Transcriptase (18090010, Invitrogen). The resulting cDNA was treated with ExoSAP-IT Reagent (78201, Applied Biosystems) at 37°C for 15 minutes, followed by incubation with 20mM EDTA and 0.1M sodium hydroxide at 70°C for 12 minutes. Hydrochloric acid (0.1M) was added to the sample to quench the reaction. The purified cDNA was isolated using Dynabeads MyONE Silane (37002D, ThermoFisher Scientific) according to manufacturer’s instructions. Briefly, 10μL of beads were magnetically separated and washed once with 500μL RLT Buffer, resuspended in 93μL RLT Buffer, and added to the samples. The bead mixture was combined with 111.6μL 100% ethanol, pipette mixed, and incubated at room temperature for 5 minutes. The sample was placed on a magnet, the supernatant was removed, and then washed twice with 1mL 80% ethanol, waiting 30 seconds between each magnetic separation. After the final wash, the beads were air dried for 5 minutes, resuspended in 5μL Tris-HCl (5mM, pH 7.5) and incubated at room temperature for 5 minutes.

#### 5’ cDNA Adapter Ligation

The sample was combined with 0.8μL rand3Tr3 oligonucleotide adaptor and 1μL 100% DMSO, incubated at 75° for 2 minutes, and then placed on ice for 1 minute. Ligation Master Mix (9μL 50% PEG 8000, 2μL 10x NEB T4 RNA Ligase Buffer, 1.1μL nuclease free water, 0.2μL 1mM ATP, 1.5μL T4 RNA Ligase I) was added to the sample, mixed at 1200 rpm for 30 seconds, and then incubated at room temperature overnight.

#### cDNA Isolation / qPCR Quantification

The cDNA was isolated using Dynabeads MyONE Silane following the instructions already described (5μL of beads per sample were used, washed with 500μL RLT Buffer, and resuspended in 60μL RLT Buffer and an equal volume of 100% ethanol). The samples were eluted in 25μL Tris-HCl (10mM, pH 7.5). A 1:10 dilution of cDNA was used to quantify the sample by qPCR. Based on the resulting Ct values, a PCR reaction was run on the diluted sample using 25μL Q5 Hot Start PCR Master Mix (M0494S, NEB) and 2.5μL (20μM) each of 2 indexed primers (Illumina TruSeq Combinatorial Dual (CD) index adapters, formerly known as TruSeq HT). The sample was amplified using a range of cycles based on the Cq obtained from the qPCR (Cq-3, Cq, Cq+3) and then visualized on a 12% TBE polyacrylamide gel to determine the optimal amount of amplification for the final library (ideally a cycle number is chosen where the amplicon has just become visible) (Supplemental Figure 2).

#### Library Amplification / Gel Purification

The undiluted cDNA library was amplified by combining 12.5μL of the sample with 25μL Q5 Hot Start PCR Master Mix and 2.5μL (20μM) of the same indexed primers used previously (amplification for the full undiluted sample will be 3 cycles less than the cycle selected from the diluted sample). The PCR reaction was isolated using HighPrep PCR Clean-up System (AC-60050, MAGBIO) according to manufacturer’s instructions. The final sequencing library was gel purified by combining the sample with 10x OrangeG DNA loading buffer and running on a 3% quick dissolve agarose gel containing SYBR Safe Dye (1:10,000). Following gel electrophoresis, a long wave UV lamp was used to extract DNA fragments from the gel ranging from 175 to 300 base pairs. The DNA was isolated using QiaQuick MinElute Gel Extraction Kit (28604, Qiagen). The purified sequencing library was analyzed via TapeStation using DNA ScreenTape (either D1000 or HS D1000) according to the manufacturer’s instructions to assess for appropriate size and concentration (the final library should be between 175 and 300 base pairs with an ideal concentration of at least 10nM).

### Overview of Snakemake Workflow

Sequencing of the cDNA libraries was primarily performed using an Illumina NovaSEQ 6000 to generate 2×150bp paired-end runs consisting of 40 million raw reads per sample (as frequency of conversions can be directly impacted by sequencing depth, we recommend a minimum of 40 million reads). The resulting reads are analyzed via a modified computational pipeline based on the original eCLIP strategy that has been converted into a Snakemake workflow (accessible via GitHub at https://github.com/ajlabuc/meCLIP). It can be executed according to Snakemake guidelines using a configuration file detailing the location of the respective sequencing files and relevant genomes. Specific commands within the pipeline are as follows: the reads are initially inspected for appropriate quality using FastQC (v. 0.11.7) and the in-line unique molecular identifier (UMI) located within the ssDNA adapter (rand3Tr3) at the beginning of read 2 is extracted using UMI-tools (v. 1.0.0) to prepare the reads for downstream de-duplication. The remaining non-random ssDNA adapter and indexed RNA adapters are then removed using Cutadapt (v. 2.4), with any reads less than 18bp being discarded. The trimmed reads are then briefly analyzed visually once more with FastQC to ensure all adapters are successfully removed. Two mapping steps are then performed using the splicing-aware RNA aligner STAR (v. 2.7.1a). First the reads are mapped to the species appropriate version of RepBase (v18.05) with any successfully mapped reads being removed from further analysis (this step ultimately leads to elimination of reads mapping to ribosomal RNA and other annotated repetitive sequences; however, if m^6^A identification within these loci are of interest then this filtering step can be turned off within the workflow and given sufficient read length many repeats should be able to be uniquely mapped). The remaining reads are then mapped to the full human genome (hg19) with only uniquely mapping reads being included in final alignment. Subsequent removal of PCR duplicates is performed with UMI-tools using the previously extracted UMIs, with the allowed error rate within the UMI itself determined by the default settings. The final alignment file is sorted and indexed and then used as input for a custom m^6^A identification algorithm (in keeping with the initial eCLIP pipeline, only read 2 is used).

### m^6^A Identification Algorithm

Putative m^6^A residues are identified using a custom analysis pipeline that utilizes the ‘mpileup’ command of SAMtools (v. 1.9) to identify variations from the reference genome at single-nucleotide resolution across the entire genome. An internally developed Java package is then employed to identify C-to-T mutations occurring 1) within the m^6^A consensus motif ‘RAC’: ‘R’ is any purine, A or G; A being the methylated adenosine; and C where the mutation occurs; and 2) within a set frequency threshold of greater than or equal to 2.5% and less than or equal to 50% of the total reads at a given position (with a minimum of 3 C-to-T mutations at a single site). The broader consensus motif ‘DRACH’, where ‘D’ denotes A, G or U, and ‘H’ denotes A, C or U can also be used for greater selectivity by modifying the configuration file. The resulting m^6^A sites are then automatically compared to those identified in the corresponding input sample and any sites occurring in both are removed from the final list of m^6^As (this eliminates any mutations that are not directly induced from the anti-m^6^A antibody crosslinking).

*Note – Previous iCLIP-based m^6^A identification strategies(Linder et al. 2015) have used crosslinking-induced truncations (CITS) to further identify m^6^A sites based on the observation that reverse transcription often terminates at the RNA:antibody crosslink site. While eCLIP does maintain the ability to identify these events at single-nucleotide resolution via ligation of the ssDNA rand3Tr3 adapter to the cDNA fragments at their 3′ ends, we do not often see this event in our meCLIP strategy (possibly due to increased fidelity of the reverse transcriptase used (SuperScript IV) compared to the older version (SuperScript III) or our use of a single antibody (Abcam) compared to others (Synaptic Systems) that are more prone to induce truncations). Therefore, our identification strategy does not include truncation-based identification of m^6^As.

## Acknowledgements

We would like to thank K. Riemondy, R. Sheridan, N. Mukherjee, and M. Taliaferro for helpful comments on the manuscript. We acknowledge the University of Colorado Cancer Center Genomics Core (supported by NIH grant P30-CA46934) for technical support. This work was supported by NIH grants T32GM008730 (J.T.), T32CA190216 (A.M.P.), and R35GM119575 (A.M.J); a Department of Defense Breast Cancer Research Program Breakthrough Fellowship Award W81XWH-18-1-0023 (A.M.P), and a seed grant from the University of Colorado School of Medicine RNA Bioscience Initiative.

## Supplemental Figures

**Supplemental Figure 1.**
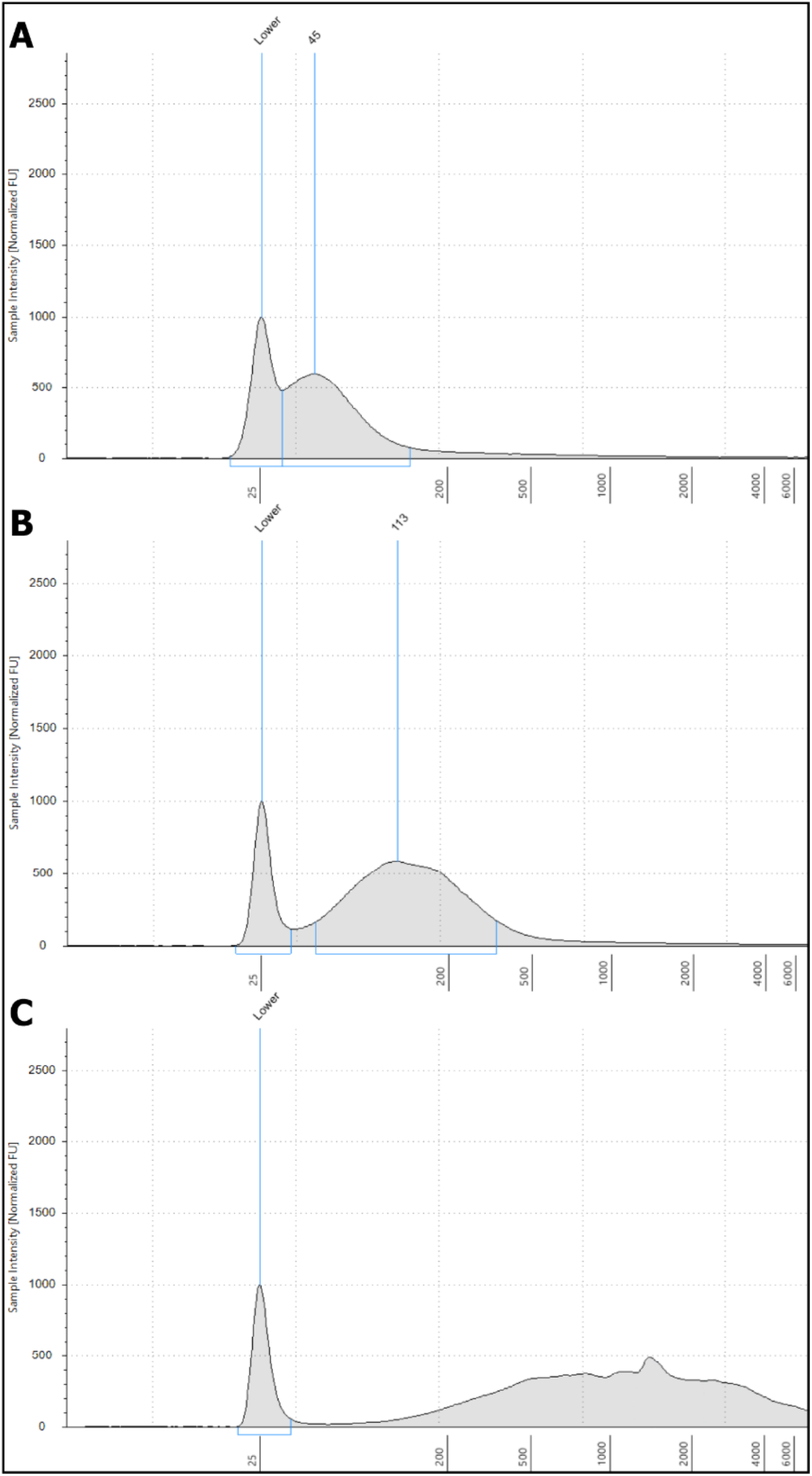
Agilent TapeStation 4200 results of RNA fragementation using High Sensitivity RNA ScreenTape. A small aliquot of total RNA (2μg) was fragemented for various durations at 70C and then a highly-diluted sample (~3ng/μL) was analyzed according to the manufacturer’s protocol. A) Example of over-fragmented sample that is outside the recommended range of 100-200bp (9 min of fragmentation). B) Optimal fragmentation result (~3 min). C) Under fragmented sample (1 min with higher concentration); although minimal under-fragmenation will likely still yield acceptable libraries, more extreme scenarios such as the one depicted will result in large amplicons that are outside the recommended range of the sequencer. “Lower” refers to the lower size marker for the TapeStation instrument.

**Supplemental Figure 2.**
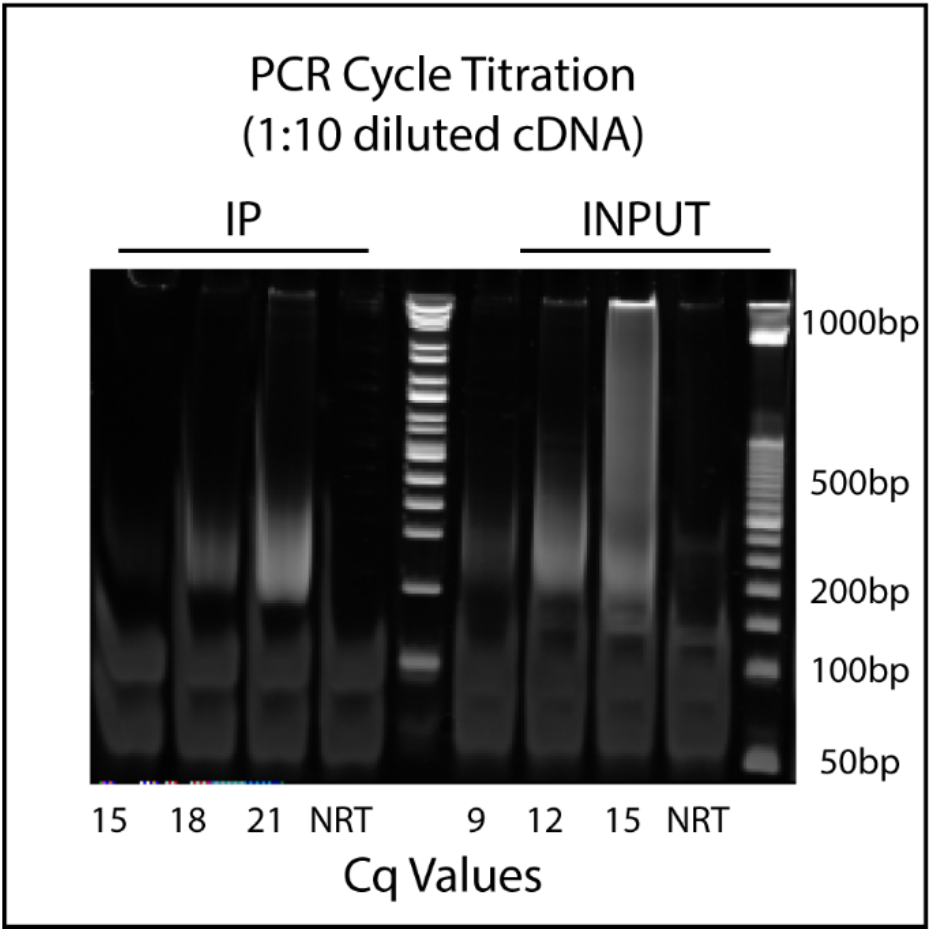
Example of PCR cycle titration used to optimize the final library amplification. Based on qPCR results obtained using a diluted sample of cDNA (1:10), the sample is amplified with three different cycles (Cq-3, Cq, Cq+3) and run on a polyacrylamide gel for visualization (i.e. the Cq obtained from qPCR for the diluted IP sample was 18, so the PCR cycle titrations were 15, 18, and 21). ‘NRT’ is a ‘no reverse transcriptase’ control. Optimal amplification typically results when the smear representing the fragment (~175-300bp) is barely visible (here 18 and 10 cycles were used for the IP and INPUT samples, respectively). Primer dimers are seen ~50-100bp. The chosen cycle is adjusted for the full library (subtract 3 cycles) and used for the final amplification.

**Supplemental Figure 3.**
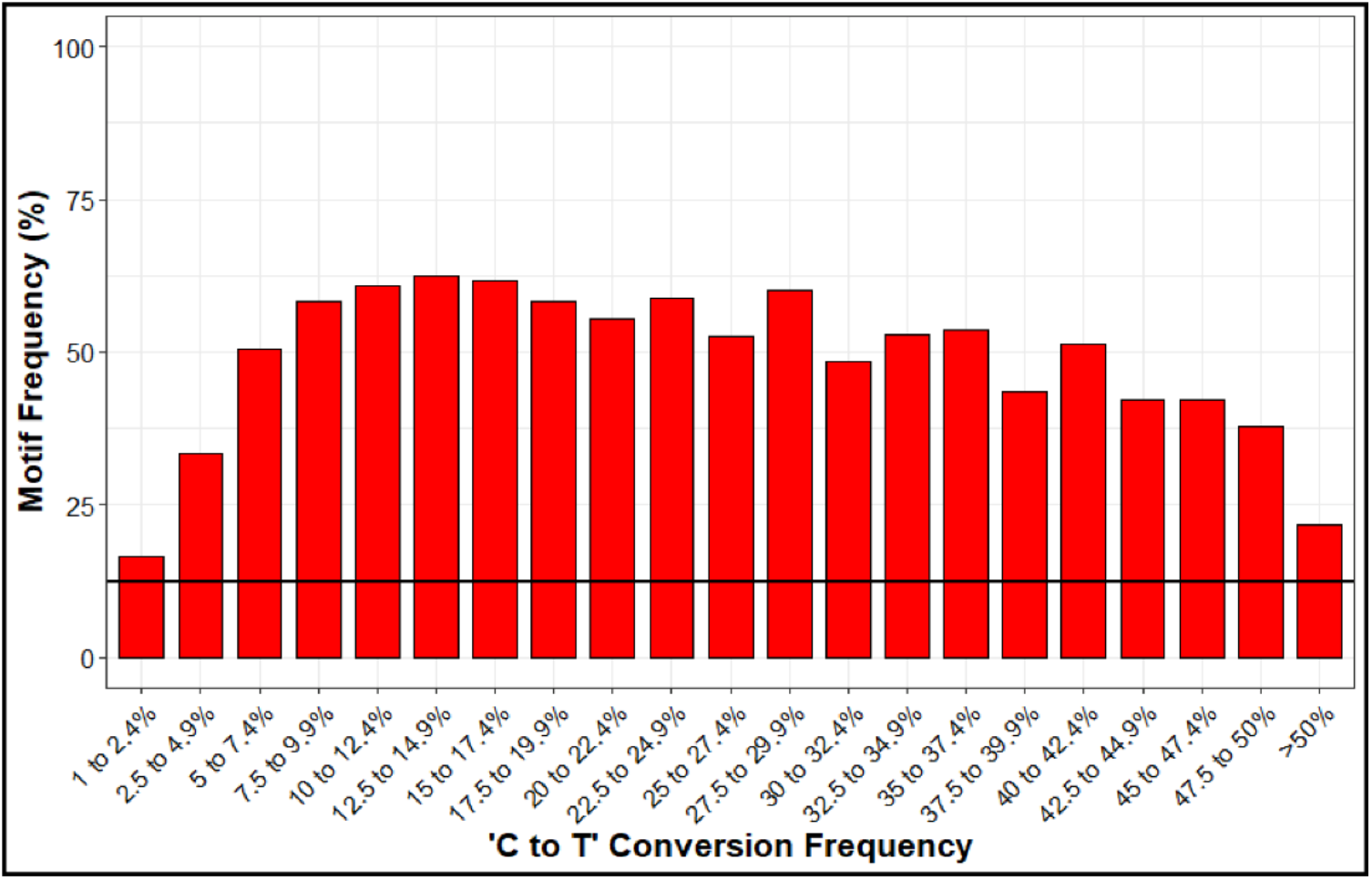
Analysis depicting the occurrence of the ‘RAC’ consensus motif relative to C-to-T mutations. Black line at 12.5% represents the random chance of having ‘RA’ upstream of the C.

**Supplemental Figure 4.**
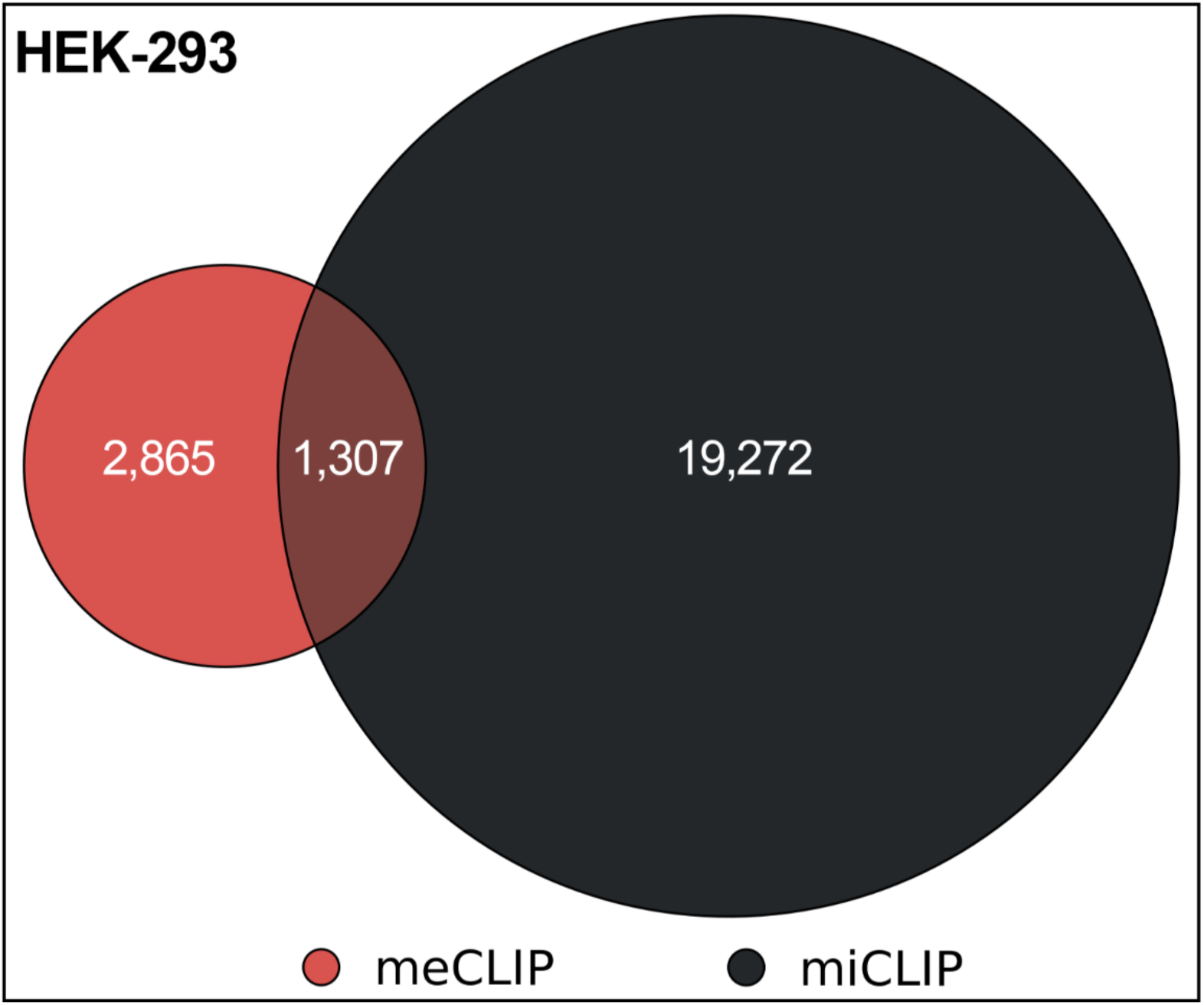
Venn diagram of m^6^A sites called in HEK-293 cells using meCLIP experimental approach and analysis pipeline compared to the m^6^A sites identified in the original miCLIP paper (Linder et al. 2015).

## References

Bao, Weidong, Kenji K. Kojima, and Oleksiy Kohany. 2015. “Repbase Update, a Database of Repetitive Elements in Eukaryotic Genomes.” Mobile DNA 6 (1): 11. https://doi.org/10.1186/s13100-015-0041-9.

Boulias, Konstantinos, D. Toczydłowska-Socha, Ben R. Hawley, Noa Liberman, Ken Takashima, Sara Zaccara, Théo Guez, et al. 2019. “Identification of the M6Am Methyltransferase PCIF1 Reveals the Location and Functions of M6Am in the Transcriptome.” Molecular Cell 75 (3): 631–643.e8. https://doi.org/10.1016/j.molcel.2019.06.006.

Chen, Kai, Zhike Lu, Xiao Wang, Ye Fu, Guan-zheng Luo, Nian Liu, Dali Han, et al. 2015. “High-Resolution N6-Methyladenosine (M6A) Map Using Photo-Crosslinking-Assisted M6A Sequencing**.” Angewandte Chemie 54: 1587–90. https://doi.org/10.1002/anie.201410647.

Dobin, Alexander, Carrie A. Davis, Felix Schlesinger, Jorg Drenkow, Chris Zaleski, Sonali Jha, Philippe Batut, Mark Chaisson, and Thomas R. Gingeras. 2013. “STAR: Ultrafast Universal RNA-Seq Aligner.” Bioinformatics 29 (1): 15–21. https://doi.org/10.1093/bioinformatics/bts635.

Dominissini, Dan, Sharon Moshitch-Moshkovitz, Schraga Schwartz, Mali Salmon-Divon, Lior Ungar, Sivan Osenberg, Karen Cesarkas, et al. 2012. “Topology of the Human and Mouse M6A RNA Methylomes Revealed by M6A-Seq.” Nature 485 (7397): 201–6. https://doi.org/10.1038/nature11112.

Doxtader, Katelyn A., Ping Wang, Anna M. Scarborough, Dahee Seo, Nicholas K. Conrad, and Yunsun Nam. 2018. “Structural Basis for Regulation of METTL16, an S-Adenosylmethionine Homeostasis Factor.” Molecular Cell 71 (6): 1001–1011.e4. https://doi.org/10.1016/j.molcel.2018.07.025.

Garcia-Campos, Miguel Angel, Sarit Edelheit, Ursula Toth, Modi Safra, Ran Shachar, Sergey Viukov, Roni Winkler, et al. 2019. “Deciphering the ‘M6A Code’ via Antibody-Independent Quantitative Profiling.” Cell 178 (3): 731–747.e16. https://doi.org/10.1016/j.cell.2019.06.013.

Geula, Shay, Sharon Moshitch-Moshkovitz, Dan Dominissini, Abed Al Fatah Mansour, Nitzan Kol, Mali Salmon-Divon, Vera Hershkovitz, et al. 2015. “M6A MRNA Methylation Facilitates Resolution of Naïve Pluripotency toward Differentiation.” Science. https://doi.org/10.1126/science.1261417.

Gilbert, Wendy V., Tristan A. Bell, and Cassandra Schaening. 2016. “Messenger RNA Modifications: Form, Distribution, and Function.” Science. https://doi.org/10.1126/science.aad8711.

Hafner, Markus, Markus Landthaler, Lukas Burger, Mohsen Khorshid, Jean Hausser, Philipp Berninger, Andrea Rothballer, et al. 2010. “Transcriptome-Wide Identification of RNA-Binding Protein and MicroRNA Target Sites by PAR-CLIP.” Cell 141 (1): 129–41. https://doi.org/10.1016/j.cell.2010.03.009.

Imanishi, Miki, Shogo Tsuji, Akiyo Suda, and Shiroh Futaki. 2017. “Detection of: N 6-Methyladenosine Based on the Methyl-Sensitivity of MazF RNA Endonuclease.” Chemical Communications. https://doi.org/10.1039/c7cc07699a.

Ke, Shengdong, Endalkachew A. Alemu, Claudia Mertens, Emily Conn Gantman, John J. Fak, Aldo Mele, Bhagwattie Haripal, et al. 2015. “A Majority of M6A Residues Are in the Last Exons, Allowing the Potential for 3ʹ UTR Regulation.” Genes and Development, 2013–15. https://doi.org/10.1101/gad.269415.115.9.

Ke, Shengdong, Amy Pandya-Jones, Yuhki Saito, John J. Fak, Cathrine Broberg Vågbø, Shay Geula, Jacob H. Hanna, Douglas L. Black, James E. Darnell, and Robert B. Darnell. 2017. “M6A MRNA Modifications Are Deposited in Nascent Pre-MRNA and Are Not Required for Splicing but Do Specify Cytoplasmic Turnover.” Genes and Development 31 (10): 990–1006. https://doi.org/10.1101/gad.301036.117.

Kong, Weili, Efraín E. Rivera-Serrano, Jason A. Neidleman, and Jian Zhu. 2019. “Single-Base Mapping of M6A by an Antibody-Independent Method.” Science Advances 5 (July): 6. https://doi.org/10.1016/j.jmb.2019.09.021.

Köster, Johannes, and Sven Rahmann. 2012. “Snakemake-a Scalable Bioinformatics Workflow Engine.” Bioinformatics 28 (19): 2520–22. https://doi.org/10.1093/bioinformatics/bts480.

Li, Heng, and Richard Durbin. 2009. “Fast and Accurate Short Read Alignment with Burrows-Wheeler Transform.” Bioinformatics 25 (14): 1754–60. https://doi.org/10.1093/bioinformatics/btp324.

Li, Heng, Bob Handsaker, Alec Wysoker, Tim Fennell, Jue Ruan, Nils Homer, Gabor Marth, Goncalo Abecasis, and Richard Durbin. 2009. “The Sequence Alignment/Map Format and SAMtools.” Bioinformatics 25 (16): 2078–79. https://doi.org/10.1093/bioinformatics/btp352.

Licatalosi, Donny D., Aldo Mele, John J. Fak, Jernej Ule, Melis Kayikci, Sung Wook Chi, Tyson A. Clark, et al. 2008. “HITS-CLIP Yields Genome-Wide Insights into Brain Alternative RNA Processing.” Nature 456 (7221): 464–69. https://doi.org/10.1038/nature07488.

Linder, Bastian, Anya V. Grozhik, Anthony O. Olarerin-George, Cem Meydan, Christopher E. Mason, and Samie R. Jaffrey. 2015. “Single-Nucleotide-Resolution Mapping of M6A and M6Am throughout the Transcriptome.” Nature Methods 12 (8): 767–72. https://doi.org/10.1038/nmeth.3453.

Liu, Jianzhao, Yanan Yue, Dali Han, Xiao Wang, Y. Fu, Liang Zhang, Guifang Jia, et al. 2014. “A METTL3-METTL14 Complex Mediates Mammalian Nuclear RNA N6-Adenosine Methylation.” Nature Chemical Biology 10 (2): 93–95. https://doi.org/10.1038/nchembio.1432.

Lorenz, Daniel A, Shashank Sathe, Jacyln M Einstein, and Gene W Yeo. 2019. “Direct RNA Sequencing Enables M6A Detection in Endogenous Transcript Isoforms at Base Specific Resolution.” RNA, rna.072785.119. https://doi.org/10.1261/rna.072785.119.

Meyer, Kate D. 2019. “DART-Seq: An Antibody-Free Method for Global M6A Detection.” Nature Methods 16 (December). https://doi.org/10.1038/s41592-019-0570-0.

Meyer, Kate D., Yogesh Saletore, Paul Zumbo, Olivier Elemento, Christopher E. Mason, and Samie R. Jaffrey. 2012. “Comprehensive Analysis of MRNA Methylation Reveals Enrichment in 3′ UTRs and near Stop Codons.” Cell 149 (7): 1635–46. https://doi.org/10.1016/j.cell.2012.05.003.

Nostrand, Eric L Van, Gabriel A Pratt, Alexander A Shishkin, Chelsea Gelboin-Burkhart, Mark Y Fang, Balaji Sundararaman, Steven M Blue, et al. 2016. “Robust Transcriptome-Wide Discovery of RNA-Binding Protein Binding Sites with Enhanced CLIP (ECLIP).” Nature Methods 13 (November 2015): 1–9. https://doi.org/10.1038/nmeth.3810.

Olarerin-George, Anthony O., and Samie R. Jaffrey. 2017. “MetaPlotR: A Perl/R Pipeline for Plotting Metagenes of Nucleotide Modifications and Other Transcriptomic Sites.” Bioinformatics 33 (10): 1563–64. https://doi.org/10.1093/bioinformatics/btx002.

Patil, Deepak P., Chun-kan Kan Chen, Brian F. Pickering, Amy Chow, Constanza Jackson, Mitchell Guttman, Samie R. Jaffrey, and Biological Engineering. 2016. “M6A RNA Methylation Promotes XIST-Mediated Transcriptional Repression.” Nature 537 (7620): 369–73. https://doi.org/10.1038/nature19342.

Price, Alexander M, Katharina E Hayer, Alexa B R McIntyre, Nandan S Gokhale, Ashley N Della Fera, Christopher E Mason, Stacy M Horner, Angus C Wilson, Daniel P Depledge, and Matthew D Weitzman. 2019. “Direct RNA Sequencing Reveals M6A Modifications on Adenovirus RNA Are Necessary for Efficient Splicing.” BioRxiv, January, 865485. https://doi.org/10.1101/865485.

Roundtree, Ian A, Guan Zheng Luo, Zijie Zhang, Xiao Wang, Tao Zhou, Yiquang Cui, Jiahao Sha, et al. 2017. “YTHDC1 Mediates Nuclear Export of N6-Methyladenosine Methylated MRNAs.” ELife 6: 1–28. https://doi.org/10.7554/eLife.31311.

Ruszkowska, Agnieszka, Milosz Ruszkowski, Zbigniew Dauter, and Jessica A. Brown. 2018. “Structural Insights into the RNA Methyltransferase Domain of METTL16.” Scientific Reports 8 (1): 1–13. https://doi.org/10.1038/s41598-018-23608-8.

Sendinc, E., D. Valle-Garcia, Abhinav Dhall, Hao Chen, T. Henriques, Jose Navarrete-Perea, Wanqiang Sheng, Steven P. Gygi, K. Adelman, and Yang Shi. 2019. “PCIF1 Catalyzes M6Am MRNA Methylation to Regulate Gene Expression.” Molecular Cell. https://doi.org/10.1016/j.molcel.2019.05.030.

Shah, Ankeeta, Yingzhi Qian, Sebastien M. Weyn-Vanhentenryck, and Chaolin Zhang. 2017. “CLIP Tool Kit (CTK): A Flexible and Robust Pipeline to Analyze CLIP Sequencing Data.” Bioinformatics 33 (4): 566–67. https://doi.org/10.1093/bioinformatics/btw653.

Smith, Tom, Andreas Heger, and Ian Sudbery. 2017. “UMI-Tools: Modelling Sequencing Error in Unique Molecular Identifiers to Improve Quantification.” Cold Spring Harbor Laboratory Press 27 (3): 491–99. https://doi.org/10.1101/gr.209601.116.

Wang, Xiao, Zhike Lu, Adrian Gomez, Gary C. Hon, Yanan Yue, Dali Han, Ye Fu, et al. 2014. “N6-Methyladenosine-Dependent Regulation of Messenger RNA Stability.” Nature 505 (7481): 117–20. https://doi.org/10.1038/nature12730.

Wang, Yang, Yue Li, Julia I. Toth, Matthew D. Petroski, Zhaolei Zhang, and Jing Crystal Zhao. 2014. “N6-Methyladenosine Modification Destabilizes Developmental Regulators in Embryonic Stem Cells.” Nature Cell Biology. https://doi.org/10.1038/ncb2902.

Wu, Baixing, Li Li, Ying Huang, Jinbiao Ma, and Jinrong Min. 2017. “Readers, Writers and Erasers of N6-Methylated Adenosine Modification.” Current Opinion in Structural Biology 47: 67–76. https://doi.org/10.1016/j.sbi.2017.05.011.

Xiao, Wen, Samir Adhikari, Ujwal Dahal, Yu-Sheng Sheng Chen, Ya-Juan Juan Hao, Bao-Fa Fa Sun, Hui-Ying Ying Sun, et al. 2016. “Nuclear M6A Reader YTHDC1 Regulates MRNA Splicing.” Molecular Cell 61 (4): 507–19. https://doi.org/10.1016/j.molcel.2016.01.012.

Zeng, Yong, Shiyan Wang, Shanshan Gao, Fraser Soares, Musadeqque Ahmed, Haiyang Guo, Miranda Wang, et al. 2018. “Refined RIP-Seq Protocol for Epitranscriptome Analysis with Low Input Materials.” PLOS Biology 16 (9): 1–20. https://doi.org/10.1371/journal.pbio.2006092.

Zhao, Boxuan Simen, Ian A Roundtree, and Chuan He. 2016. “Post-Transcriptional Gene Regulation by MRNA Modifications.” Nature Reviews Molecular Cell Biology. Nature Publishing Group. https://doi.org/10.1038/nrm.2016.132.

